# Genetic determinants of antibiotic resistance and the evolution of trade-offs during adaptation in a single patient

**DOI:** 10.1101/2021.10.06.463321

**Authors:** Robert J. Woods, Camilo Barbosa, Laura Koepping, Juan A. Raygoza Garay, Michael Mwangi, Andrew F. Read

## Abstract

The processes by which pathogens evolve within single hosts dictate the efficacy of treatment strategies designed to slow antibiotic resistance evolution and influence the population-wide resistance levels. The aim of this study is to describe the underlying genetic and phenotypic changes leading to antibiotic resistance within a single patient who died as resistance evolved to available antibiotics. We assess whether robust patterns of collateral sensitivity and response to combinations exist that might have been leveraged to improve therapy. Whole-genome sequencing was completed for nine isolates taken from this patient over 279 days of chronic infection with *Enterobacter hormaechei*, along with systematic measurements of changes in resistance against five of the most relevant drugs considered for treatment. The entirety of the genetic change is consistent with *de novo* mutations and plasmid loss events, without the acquisition of foreign genetic material via horizontal gene transfer. The isolates formed three genetically distinct lineages, with early evolutionary trajectories being supplanted by previously unobserved multi-step evolutionary trajectories. Importantly, no single isolate evolved resistance to all of the antibiotics considered for treatment against *E. hormaechei* (*i*.*e*., none was pan-resistant). Patterns of collateral sensitivity and response to combination therapy revealed contrasting patterns across this diversifying population. Translating antibiotic resistance management strategies from theoretical and laboratory data to clinical situations, such as this, may require managing diverse populations with unpredictable resistance trajectories.

## Introduction

The effective treatment of bacterial infections has transformed modern medicine, but the ability of bacteria to evolve resistance to essentially all antibiotics now threatens these gains (CDC, 2019). For certain infections, the evolution of resistance within individual patients undergoing therapy can lead directly to treatment failure and worse patient outcomes (Folkesson et al., 2012; Gumbo et al., 2014a; Hansen et al., 2012; Woods and Read, 2015). Understanding how pathogens adapt within patients during these infections is crucial to identifying treatment strategies that better impede resistance evolution and prevent possible transmission to others (Mideo et al., 2008; zur Wiesch et al., 2011). For instance, within-host competition can modulate the evolution of drug resistance and pathogenicity, ultimately dictating which genotypes dominate the infection and which are transmitted between-hosts (Huijben et al., 2018; Mideo et al., 2008). Furthermore, understanding the genetic mechanisms under selection within a single host can help answer questions about the complex population structures resulting from chronic infections like those seen in patients with cystic fibrosis with *Pseudomonas aeruginosa* (Marvig et al., 2015b, 2015a). However, the prevailing low levels of predictability of resistance evolution at the single patient level are a major challenge to physicians on a daily basis.

A previous case study reported a patient suffering from a multispecies infection for more than 500 days that could not be resolved, resulting in the death of the patient as drug resistance progressed (Woods and Read, 2015). The infection developed in two phases (Figure 1A): An initial period dominated by methicillin-resistant *Staphylococcus aureus* (MRSA) lasting around 8 months and resulting in evolved resistance to clindamycin and daptomycin (Woods and Read, 2015). After 240 days *Enterobacter hormaechei* was cultured from the site of infection, and MRSA was never detected again. The patient succumbed to the infection 279 days after the *E. hormaechei* was first cultured and once resistance to meropenem evolved (Woods and Read, 2015). While the retrospective reconstruction of this single case cannot identify what the optimal course of treatment could have been, our goal is to better understand the evolutionary process leading to treatment failure in order to propose better strategies in the future. Specifically, we want to examine four questions focusing on the second phase of infection with *E. hormaechei*: (1) Did resistance emerge from the acquisition of *de novo* mutations or genetic elements from other bacteria? (2) Did the course of treatment select for a pan-resistant variant of *E. hormaechei* as previously described for other pathogens (Mwangi et al., 2007)? (3) Was there a systematic change in resistance against particular antibiotics leading to substantial drops in resistance against other drugs (*i*.*e*., collateral sensitivity)? And (4) were there drug combinations that could allow selection inversion approaches to be used?

**Figure 1.**
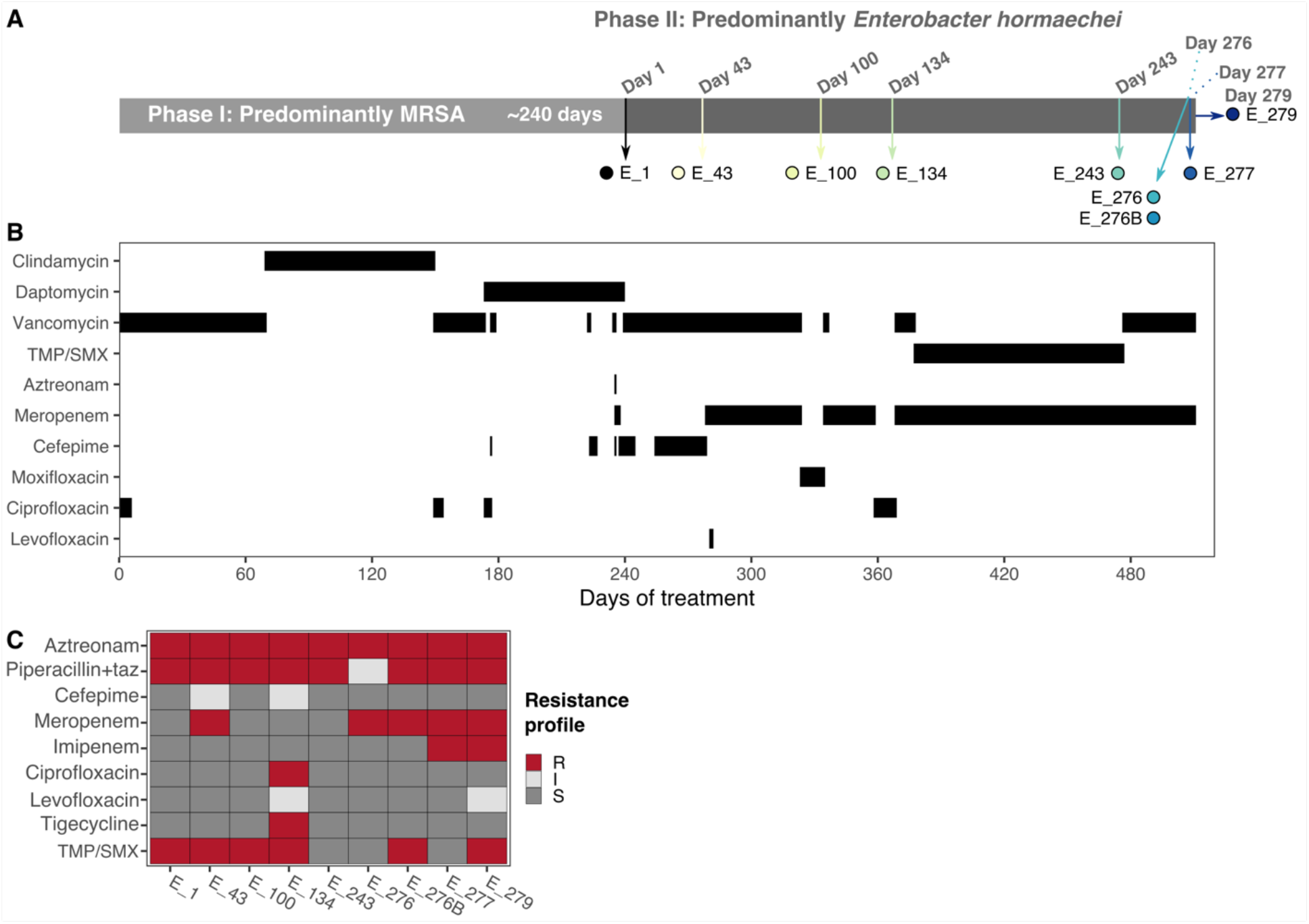
Course of infection and *Enterobacter* isolation regime. (A) The full course of infection consisted of two phases: The first infection phase (light grey) predominantly involved MRSA, lasting approximately 240 days. After that, *E. hormaechei* was detected, initiating the second phase of infection (dark grey). Phase II lasted 279 days including seven hospital visits from which nine isolates were obtained (note the points and arrows have colors maintained throughout this manuscript). The day of isolation during the second phase is indicated in dark grey at the top of the timeline. (B) Patient’s course of antibiotic treatment. (C) Resistance profile of the isolates as inferred from the CLSI breakpoints for all isolates against nine different antibiotics; resistant, intermediate and sensitive isolates are shown in red, light grey and dark grey, respectively

We addressed these questions by identifying the genetic and phenotypic changes occurring during the 279 days of evolution of *E. hormaechei* within this patient (Woods and Read, 2015). We identified the genetic changes and population diversity leading to antibiotic resistance using whole genome sequencing of the nine *E. hormaechei* isolates. We also evaluated evolutionary trade-offs in the form of collateral sensitivity, as well as changes in drug interactions across the diversifying population (Baym et al., 2016). Altogether, our data indicate that resistance evolution within this patient was entirely the product of de novo genetic changes; there was no evidence for the acquisition of foreign DNA through horizontal gene transfer. No single isolate was resistant to all possible antibiotics, instead multiple independent subpopulations accumulated multiple distinct mutations, resulting in the population exploring the antibiotic resistance genetic landscape. This multi-step evolutionary process included periods of collateral sensitivity alternating with collateral resistance. The complexity of the resulting pattern suggests that combating resistance within host will require a more comprehensive approach complemented by intensive sampling of the population within and across time points. General guidelines for clinical decision making based on such evolutionary strategies remain elusive.

## Results

We considered nine *E. hormaechei* isolates obtained during the second phase of infection within a single patient that was previously described in detail (Figure 1A; see (Woods and Read, 2015) for a full account of the clinical case description). We obtained the isolates at times when the patient visited the hospital and further analyzed them by using whole genome sequencing, estimating changes in resistance against five antibiotics and evaluating drug interactions between meropenem and four other drugs.

### Genomic analyses

To better understand the genetic basis of resistance in this patient, we performed whole genome sequencing analysis of the nine *E. hormaechei* isolates. We generated a parsimony tree by hand, resulting in no homoplastic genetic changes showing three distinct lineages with no shared variants (Figure 2A and B). The two lineages including the early isolates were not detected subsequently, so they were potentially lost during the course of treatment. Importantly, the use of vancomycin and TMP/SMX during treatment was not targeted at *E. hormaechei* but were included to reduce the risk of a MRSA re-emergence (Fig 1B). Yet, these antibiotics, particularly the TMP/SMX, could still have affected the *E. hormaechei* population structure. Indeed, the late isolates revealed loss of TMP/SMX resistance associated with loss of the plasmid encoded *dfrA12* and sulphonamide resistance *sul1* genes (Figure 1C and 2A). Overall, the final population was genetically diverse with more than half of the observed genetic variation still present during the last three days of the infection.

**Figure 2.**
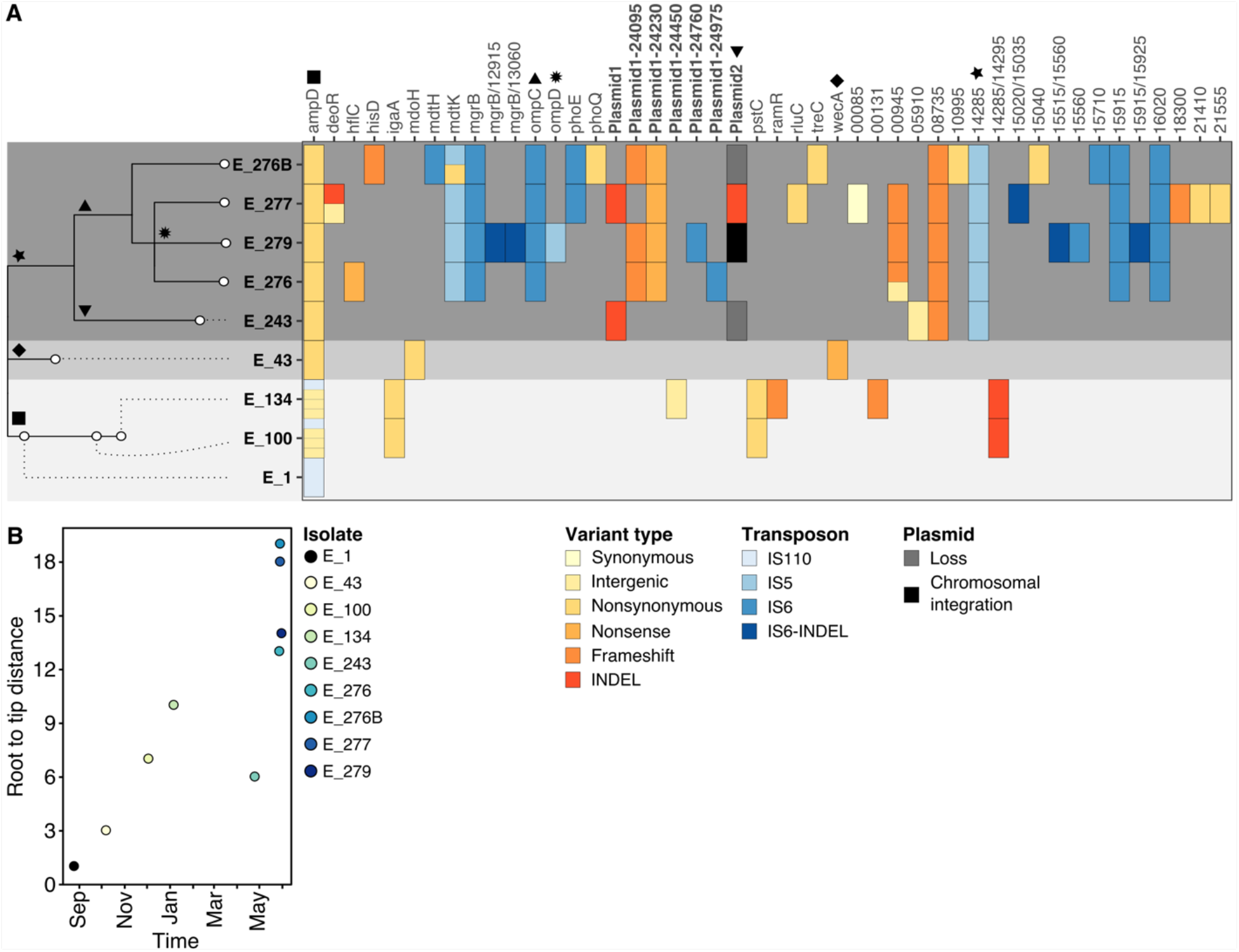
Genomics of adaptation within a single patient. (A) Maximum parsimony phylogeny of the nine *E. hormaechei* isolates taken from a single patient and heatmap of all the 58 variants in 40 genes of the nine isolates. Three major lineages were observed with no shared variants after day 243 (E_243) of infection with *E. hormaechei* are highlighted in grey blocks. The isolates recovered in the last four days harbor more than half of the genetic diversity observed during the entire period. The variant type in the heatmap is indicated in different shades of red, transposon activity is indicated in different shades of blue and plasmid loss or integration is shown in dark grey and black squares. Genes in bold correspond to variants observed within plasmids. Symbols highlight genes potentially associated with changes in resistance against different antibiotics relevant for each lineage. (B) Root to tip distance for each isolate over time. The isolates are shown in different shades of blue, with the earliest isolate (E_1) shown in black. The following source data is available for Figure 2: Figure 2 – Source data 1.

We found no evidence of the acquisition of foreign genetic elements in our analysis. Instead, the evolution of resistance in this context was mainly driven by the accumulation of genetic change due to insertion sequence (IS) activity, small INDELS and point mutations. Notably, there was no single isolate across the different lineages accumulating mutations and phenotypic resistance to all of the drugs considered for treatment (Figure 1C and 2A). We identified a total of 58 mutations across all the isolates in 39 chromosomal genes and 5 genes encoded in one of the three plasmids found on the earliest isolate (Figure 2A and Figure 2 Source data 1). A large fraction of all the identified variants were the result of transposon activity or plasmid loss (Figure 2A): 15 insertions, deletions or rearrangements of three different insertion sequence elements (IS6-like, IS5-like and IS10-like), five large deletions (1.7-34kb) adjacent to a copy of an IS6-like transposon, and the partial (∼70% loss) or complete loss of two of the plasmids were identified. Of all mutations, we only found a single synonymous and seven intergenic variants while most of the remaining variants were strongly disruptive (Figure 2A). This pattern strongly suggests that non-neutral evolution was predominant in this particular context.

Some of the identified mutations are associated with resistance against particular drugs that were not used during treatment, suggesting the potential evolution of collateral resistance. For instance, isolate E_134 was resistant to tigecycline (Figure 1B and C, and 2A), an antibiotic to which the patient was not exposed. This isolate had a frameshift variant in *ramR*, a TetR-like transcriptional regulator mediating the expression of *romA* and *ramA* (Figure 2A). Loss-of-function mutations in *ramR* have been associated with multidrug resistance in *Enterobacter, Klebsiella pneumoniae* and *Salmonella enterica* serovar *Typhimurium* (Majumdar et al., 2015; Ricci and Piddock, 2009; Rosenblum et al., 2011; Veleba et al., 2013). Similarly, a variant in *phoQ* was identified in one of the later isolates, E_276B (Figure 2A), a gene that has been commonly associated with resistance against colistin and some aminoglycosides (Band et al., 2016). Yet, neither of these antibiotics were given, and the isolates were not phenotypically resistant. Altogether, these data demonstrate that as this population evolved in response to the administered drugs, it also evolved heterogeneity that may impact success in future, changing environments.

Parallel evolution at the gene level was found for a single gene, *ampD*. While all isolates had at least one variant, there were three independent origins: an IS insertions 11bp upstream of *ampD* (E_1 and descendant isolates), 9 bp insertion between bp 137 and 138 (E_43), and a single nucleotide polymorphism causing amino acid change T137K (E_243 and later isolates, Figure 2A). Mutations in *ampD*, a repressor of the AmpC β-lactamase, lead to increases in resistance against numerous β-lactams in distinct bacterial pathogens (Barnaud et al., 2001; Bratu et al., 2007; Langaee et al., 2000; Naas et al., 2001). All of the isolates were indeed resistant to the commonly used β-lactams, aztreonam and piperacillin/tazobactam, but most retained sensitivity to carbapenems (Figure 2A), a pattern consistent with AmpC overexpression (Jacoby, 2009).

Other identified variants have been associated with resistance to β-lactams. Variants in *deoR, wecA*, and a large transposon mediated deletion affecting *rcsC*, which are genes involved in cell wall synthesis, O-antigen production and cell division and are commonly associated to virulence and β-lactam resistance (Henze and Berger-Bächi, 1995; Laubacher and Ades, 2008; Lehrer et al., 2007). The observed nonsense variant in *wecA* emerged early during the infectious process but was not seen later, despite potentially conferring resistance against meropenem, the main treatment agent in this case (E_43; Figure 2A and 1B). Similarly, the two variants observed late during the infection in *deoR* were not present in E_277, an isolate obtained two days later (Figure 2A).

We identified mutations in genes known to confer resistance against carbapenems, the main treatment drug used in the later stages of this case (meropenem). The four latest isolates shared a transposon insertion within *ompC* (Figure 2A), a gene coding for an outer membrane porin. Resistance against most carbapenems and some cephalosporins has been associated with decreased expression of this gene (Yigit et al., 2002). Additionally, some isolates had similar transposon events affecting *ompD* and *phoE*, a porin and a phosphoporin involved in β-lactam uptake, respectively (Chia et al., 2009; Dargent et al., 1986; Kaczmarek et al., 2006; Su et al., 2012).

Lastly, there was partial or complete loss of plasmids in most isolates within the later lineage. One of these plasmids (Plasmid 2) harbors genes associated with resistance against trimethoprim (*dfrA12*), aminoglycosides (*aadA2*), sulfonamides (*sul1*) and fluoroquinolones (*qnrS1*) (Corkill et al., 2005; Domínguez et al., 2019; Frank et al., 2007; Grape et al., 2003; Lee et al., 2001; Phillips et al., 1986). The loss and integration of this plasmid coincided with changes in sensitivity against TMP/SMX, but not against fluoroquinolones or aminoglycosides (Figure 1C) in the later isolates. Ultimately, no single clone dominated the population, suggesting that mechanisms that maintain diversity, such as niche differentiation, fitness costs of resistance or clonal interference, played an important role during the evolutionary dynamics taking place within this patient.

### Changes in resistance

Antibiotic susceptibility testing (calculation of IC_90_) was repeated to better differentiate subtle resistance evolution and correlations, either positive or negative, in resistance to the five most relevant antibiotics considered for this patient (Figure 3 and Figure 3 – Source data 1).

**Figure 3.**
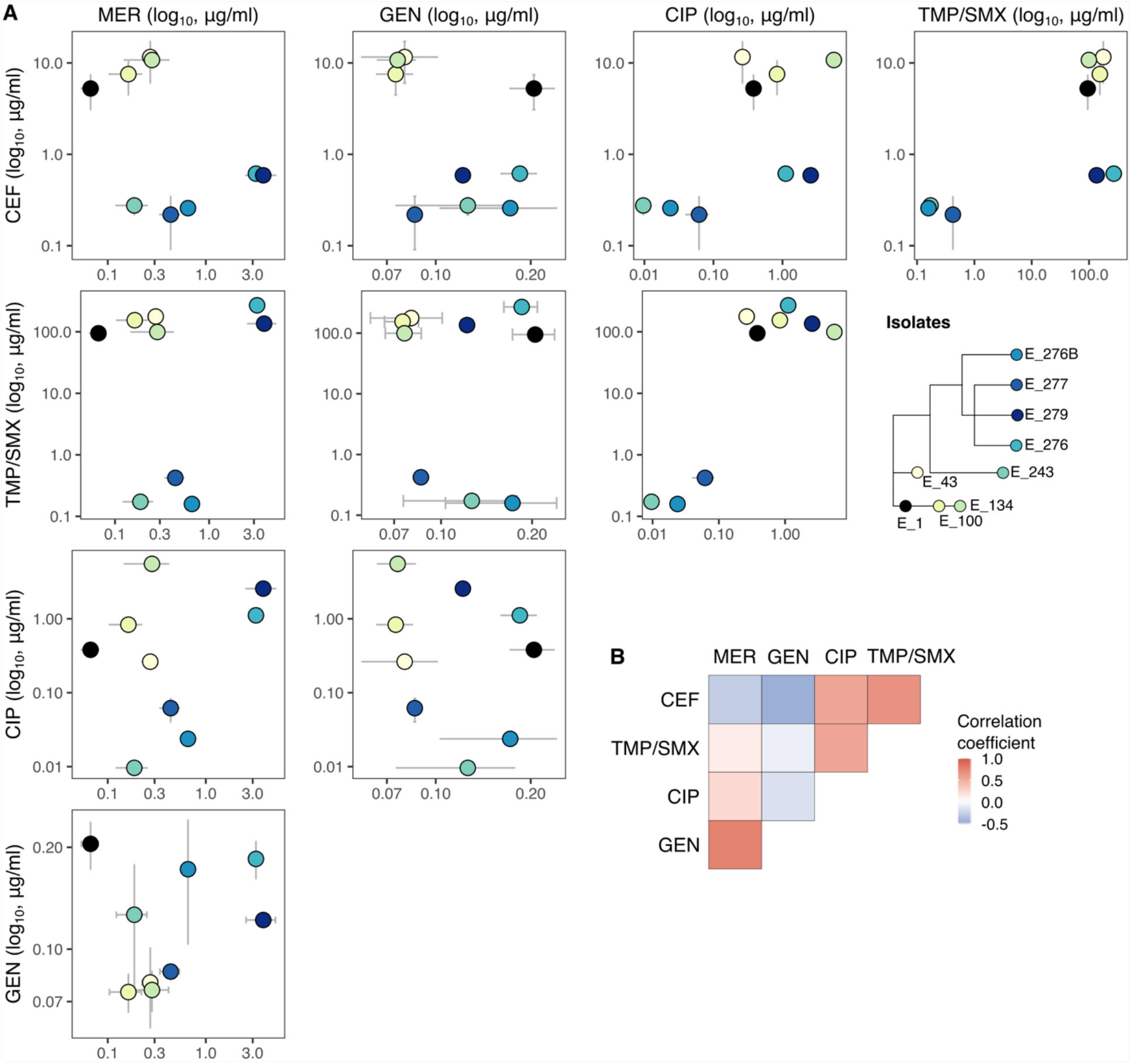
Changes in resistance against five different antibiotics. (A) The concentration inhibiting 90% of growth for meropenem (MER), cefepime (CEF), gentamicin (GEN), ciprofloxacin (CIP) and sulfamethoxazole/trimethoprim (TMP/SMX, or bactrim) is plotted for all combinations of the five antibiotics and for all isolates. The earliest isolate is shown in black and all other isolates in different shades of yellow, green and blue. Points and error bars correspond to Mean IC_90_ ± SD of three independent replicates. (B) The Spearman’s rank correlation coefficient is shown in a heatmap for all possible associations between the five drugs (−0.5 < ρ_s_ < 0.78). After FDR correction for multiple testing no significant associations were found between the changes in resistance between the five drugs (*P* > 0.28). The following source data is available for Figure 3: Figure 3 – Source data 1.

Meropenem and cefepime predominantly showed a pattern of collateral resistance, with rare collateral sensitivity. The first four isolates, gathered when cefepime was the predominant antibiotic, showed a tendency for increased cefepime IC_90_, and importantly, collateral resistance to meropenem, despite meropenem not being used (Figure 3A; top left panel). The later five isolates, obtained when treatment had predominantly shifted to meropenem, also showed positive correlation between meropenem and cefepime IC_90_. However, comparing the early group to the later one, resistance against cefepime dropped several orders of magnitude, while sensitivity against meropenem remained at comparable or slightly higher levels (Figure 3A; top panel). These early and later groups are separated by a single branch on the parsimony tree that contains three genetic changes. These mutations are the aforementioned mutation in AmpD, T137K, a transposon insertion into CPT31_14285 a predicted glycosyltransferase, and a frame shift in CPT31_08735, a TetR/AcrR family transcriptional regulator. Often, a TetR-like regulator represses the transcription of the divergently transcribed gene (Ramos et al., 2005). The divergently transcribed gene here is CPT31_08740, which has homology to the multiple stress resistance protein BhsA. The meropenem resistant isolates from the end of the infectious process also had higher levels of resistance against cefepime, yet they never reached the levels of resistance of the early group.

Similarly, the pairwise correlations across all nine isolates for all combinations of these five antibiotics revealed both positive and negative correlations (Figure 3A and B). However, none of these associations were statistically significant (−0.5 < ρ_s_ < 0.78, *P* > 0.28; Figure 3B), and careful inspection of the genetic changes reveal a more complex picture. First, sensitivity against gentamicin remained below the initial levels of resistance of the earliest isolate, suggesting that increases in resistance against any other drug did not co-select for resistance against this drug. Second, the drastic drop in resistance against ciprofloxacin and TMP/SMX coincides with the loss of plasmid 2 in some of the populations in the late lineage, which carries resistance against these drugs. Lastly, none of the isolates showed phenotypic resistance against all of the five antibiotics tested; isolate E_134 had the most resistances, to three of the five antibiotics tested (CIP, TMP/SMX and CEF).

### Antibiotic combinations

Drug combinations can be used to invert selection when mutations that increase resistance to one drug result in higher sensitivity against a particular combination of antibiotics (Baym et al., 2016). We systematically evaluated each drug when combined with meropenem by plotting growth at various combinations in so-called checkerboard plots (Figure 4 and Figure 4 – Source data 1). We first consider combinations with cefepime (Figure 4A and Figure 4 – Supplementary Figure 1 and 2). Evolution among the early isolates reshape the growth surface area by expanding growth into higher concentration of both drugs, such that clones with increased resistance to meropenem also have increased resistance to cefepime at all concentrations (Isolates E_1, E_43, E_100 and E_134; Figure 4A). However, in the later five isolates, there evolved a one-directional inhibition of antibiotic efficacy, whereby small amounts of meropenem allow *E. hormaechei* to grow in much higher concentrations of cefepime. Among these later five isolate the shape is similar, but there is a re-scaling along both the cefepime and meropenem axes. Thus, the opportunity for selection inversion only exists between these two clades, but within each clade there is a consistent pattern of collateral resistance. The patterns observed with the remaining three drugs similarly reveal that the majority of the phenotypic evolution is stretching or shrinking along one or the other axis, and therefore do not identify drug combinations that would allow for selection inversion (Figure 4B-D). There are subtle shape changes, which correspond to the loss of plasmid 2 in some of the late isolates.

**Figure 4.**
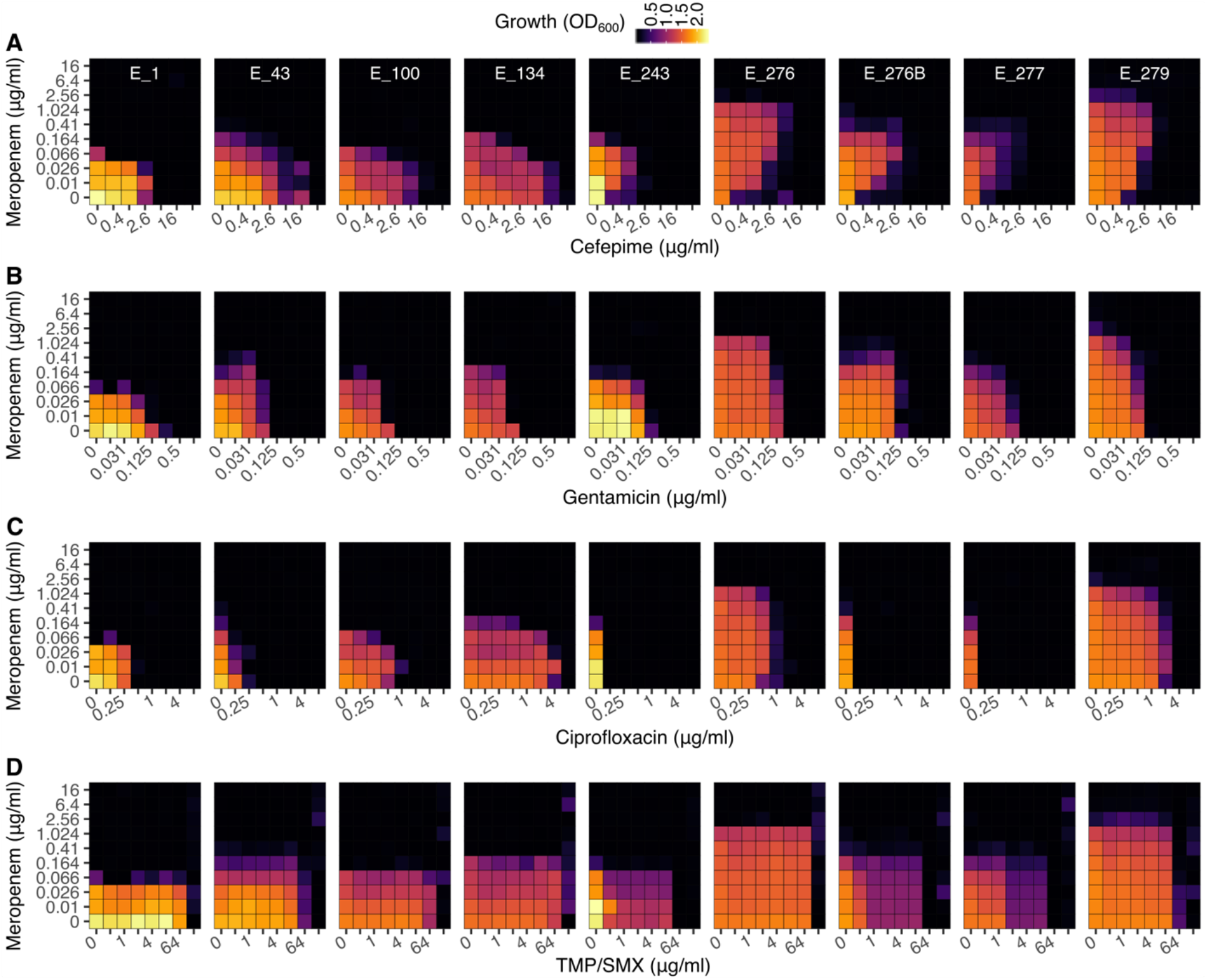
Antibiotic interactions. We measured growth in a grid of increasing concentrations of meropenem (Y-axis) and (A) cefepime, (B) gentamicin, (C) ciprofloxacin and (D) sulfamethoxazole/trimethoprim (TMP/SMX, or bactrim; X-axis) using the checkerboard assay. We measured optical density after 21 h of incubation at 36°C in two independent replicates for each isolate and drug combination showing the same broad patterns (Figure 4 – Supplementary Figure 2). Each panel corresponds to the mean optical density at each drug combination for each isolate. Areas of no growth are shown in black, with growth highlighted by the different shades of purple and yellow. The following source data and supplementary figures are available for Figure 4: Figure 4 – Source data 1, and Figure 4 – Supplementary Figure 1 and 2.

## Discussion

We systematically described the evolutionary history of *E. hormaechei* culminating in multidrug resistance within a single patient. We found that resistance was the result of the accumulation of mutational events – rather than through horizontal acquisition of resistance genes, or invasion of a separate strain – resulting in a difficult-to-treat genetically diverse population. We identified three lineages in our analysis with important genetic and phenotypic differences. One lineage was observed early during the infection, showing steady increments in resistance against multiple antibiotics including cefepime and meropenem, which were the main treatment choices at the time. However, this early lineage was not seen again, apparently overtaken by a genetically distinct lineage that emerged between days 134 and 243 of infection with *E. hormaechei*. This later lineage had increasing resistance to meropenem but decreased resistance to cefepime. The transition between the early and late lineages represents an evolutionary trade-off that could potentially be exploited to push the population toward increased sensitivity against cefepime, so the cefepime could be used therapeutically again. However, whether leveraging this trade-off is a viable treatment strategy is unknown since evidence of the effectiveness of exploiting collateral sensitivity within a single patient remains elusive. In fact, these samples hint at the danger of such a strategy: while the between-clade pattern shows collateral sensitivities, the within-clade patterns support collateral resistance.

The genetic evolution leading to resistance within this patient was complex. Resistance was a multi-step evolutionary process, as we see at least three distinct lineages accumulating multiple mutations. With the exception of *ampD*, the resistance was all in different genes, indicating that the population was able to explore the fitness landscape by taking multiple steps in multiple directions. Moreover, the bulk of the genetic changes were the result of plasmid loss, repeated transposon activity, or mutations that were disruptive to gene function. This pattern is in line with previous findings in genomic epidemiology that have shown resistance frequently evolves *de novo* within individuals, but only a subset of clones are easily transmissible and so go on to further accumulate resistance by means of horizontal gene transfer (Baker et al., 2018; MacLean and Millan, 2019). The observed patterns of gene disruption, gene loss, and particularly the expansion of IS elements (Wagner, 2006), bring into question whether this strain could compete outside of this host.

Regardless of the underlying mechanism, the fact remains that managing and predicting this evolved diversity remains problematic for physicians at the bedside. More information on how pathogens evolve within single patients as a response to treatment is urgently needed to improve clinical decision making (Zhou et al., 2020). The present study identifies a potential window of opportunity to exploit collateral sensitivity during the transition between lineages. How to exploit this opportunity and the potential consequences of changing treatment based on this evidence remain elusive. The bulk of *in vitro* studies evaluating collateral sensitivity indicate that evolutionary trade-offs are pervasive across many species, but its predictability depends on the specific genetic mechanism of resistance being selected (Barbosa et al., 2017; Imamovic and Sommer, 2013; Lázár et al., 2014, 2013; Maltas and Wood, 2019; Nichol et al., 2019). Furthermore, the effectiveness of strategies that leverage collateral sensitivity such as cycling regimes – alternating between antibiotics with known reciprocal collateral effects – may be reduced by negative epistatic interactions between resistance mechanisms against the different drugs (Barbosa et al., 2019). Still, evidence of collateral sensitivity emerging in clinical infections is scarce, mainly because direct causation between drug use, genetic changes and changes in resistance phenotypes is difficult to demonstrate, even in longitudinal studies (Jansen et al., 2016; Imamovic et al., 2018). It is thus crucial, not only to identify patterns of collateral sensitivity among clinical isolates, but also to evaluate different modes of implementation, and potential side effects, in ways that are meaningful and informative for clinical decision making.

In the context of this patient, antimicrobial treatment predominantly involved the use of more than one drug simultaneously (Woods and Read, 2015) (Figure 1B). However, this was not motivated by the use of combination therapy targeting *E. hormaechei*. Instead, combination therapy was aimed at multispecies infection due to the potential re-emergence of the MRSA observed during the first phase of the infection (Figure 1A). Treatment typically consisted of vancomycin or TMP/SMX to fight MRSA, and a β-lactam or fluoroquinolone to treat *E. hormaechei*. Interestingly, we found that in the presence of TMP/SMX the isolates within the late lineage lost a plasmid, leading to a drop in resistance against TMP/SMX and ciprofloxacin (See ref. 7, and Figure 1 and 2). This loss of TMP/SMX resistance despite treatment being continued, points to incomplete understanding of drug penetration and selective pressure in this setting.

Combination therapy may represent the best treatment strategy against infections when resistance emerges through *de novo* mutation, as observed in this patient. The effectiveness of combination therapy stems from the potential to control the population when resistance to several drugs requires independent mutations to resist each agent, as observed in other diseases (Douglas et al., 2010; Gumbo et al., 2014b; Martin et al., 2008). However, combinations can be thwarted by collateral sensitivity to the individual drug or to their combinations. Laboratory experiments suggest that, carefully chosen combinations, that interact suppressively or synergistically with known evolved collateral sensitivity, can limit the evolution of resistance or even select against resistant cells (Barbosa et al., 2018; Chait et al., 2007; Hegreness et al., 2008; Michel et al., 2008; Torella et al., 2010). The data presented here suggests that the combination of cefepime and meropenem have a suppressive interaction, such that the most meropenem-resistant cells were best killed by cefepime in the complete absence of meropenem (see E_1, E_243 and E_277 in Figure4A).

In this chronic infection, the selected course of antimicrobials did not lead to the emergence of a single, dominant pan-resistant pathogen with accumulated resistance to all possible treatments. Instead, resistance to different key therapeutic drugs emerged among clones from the different lineages. This suggests that the use of multiple antibiotics may not consistently lead to the emergence of multidrug resistant variants, even when the community as a whole contains the multidrug resistance. This outcome contrasts with previous studies in *S. aureus*, were antibiotics progressively selected for resistance in a stepwise manner within a single clone, ultimately leading to an untreatable pan-resistant pathogen (Mwangi et al., 2007).

## Conclusions and implications

Understanding how pathogens evolve antibiotic resistance within single patients is crucial to optimizing clinical decision making, improving patient outcomes and limiting the evolution and subsequent transmission of multidrug resistance organisms. Here we described the main genetic and phenotypic changes occurring within a patient after a single run of the evolutionary tape (Gould, 1990), and identified complex evolutionary trajectories, with evolution of collateral sensitivity and resistance, and idiosyncratic changes in drug-drug interactions over time. The emerging picture reveals the challenges that arise when translating theoretically derived treatment strategies to single patients.

## Methods

### Clinical isolates and antibiotics

We recovered nine isolates from the patient during the phase when *E. hormaechei* was detected. Labels, collection dates and culture sites for each isolate are given in Table 1. We repeated MIC testing for the specific clones that underwent whole genome sequencing and phenotypic testing, which may differ from those previously reported which were based on routine clinical phenotyping (Woods and Read, 2015). We systematically measured the phenotypic changes of the isolates using five of the most relevant drugs considered during treatment of the patient, cefepime, meropenem, ciprofloxacin, gentamicin and sulfamethoxazole/trimethoprim (bactrim, hereafter abbreviated TMP/SMX). For these selected drugs, we prepared stock solutions according to manufacturer’s recommendations to be used in additional assays.

**Table 1.** Isolates names and sampling source.

| Isolate name | Day of isolation | Source of isolation |
| --- | --- | --- |
| E_1 | 1 | Blood |
| E_43 | 43 | Blood |
| E_100 | 100 | Blood |
| E_134 | 134 | Blood |
| E_243 | 243 | Wound |
| E_276 | 276 | Blood |
| E_276B | 276 | Wound |
| E_277 | 277 | Wound |
| E_279 | 279 | Blood |

### DNA extraction

We extracted genomic DNA using overnight cultures of each isolate in lysogeny broth (LB) (Bertani, 1951) and using the DNeasy Blood Tissue kit (Qiagen) following the protocol for genomic DNA Isolation from Gram-Negative Bacteria.

### Genomic analysis

Sequencing was performed at the University of Michigan Sequencing Core utilizing Illumina HiSeq and PacBio sequencing. We obtained PacBio long-read sequences for the first and last samples (E_1 and E_279), while Illumina short read sequences were obtained for all nine isolates using 100bp paired end reads and repeated for 4 isolates with 150bp reads. We assembled the genomes of the first and last isolates as follows: assemblies were generated from the PacBio data using CANU, version 1.5 (Koren et al., 2017), circularized using circlator (Hunt et al., 2015), and then polished with the obtained illumina reads. The closed genomes for strains E_1 and E_279 are available in the NCBI database (accession numbers CP023569 and CP027111 respectively). We identified genetic variants by mapping the short-read sequencing back to the closed genomes. Finally, we evaluated the presence of foreign genetic elements via horizontal gene transfer by aligning the assembled genomes of the first and last isolates using bwa (Li and Durbin, 2010). Any unmapped reads were then extracted with samtools (Li et al., 2009) and *de novo* assembled using spades (Prjibelski et al., 2020). We examined the obtained contigs and found for all isolates a ∼5kb phage phiX178 which is added as a control in the Illumina sequencing, there were typically less than 9 contigs per sample, most of the contigs were less than 250bp long and they predominantly consisted of homopolymers. This suggests that no foreign genetic material was transferred horizontally among the isolates.

### Changes in resistance

We determined the concentration inhibiting 90% of growth (IC_90_) using the broth microdilution method detailed in the CLSI standard M07 for each of the *E. hormaechei* isolates. We added each of the isolates to microdilution plates containing 2-fold dilutions of each of the antibiotics in triplicate and incubated them at 36 °C for 21 h in LB media (Bertani, 1951). At this time, we measured optical density (OD_600_) using a plate reader (FLUOstar Omega from BMG Labtech). We then fitted Hill-Curves to the OD_600_ data to calculate the IC_90_ using the R platform (R Core Team, n.d.) and the minpack.lm library (Elzhov et al., 2015).

### Checkerboard and isobologram assay

To evaluate susceptibility of the isolates to antibiotic combinations we used checkerboard assays. We diluted increasing concentrations of any two drugs along the X- and Y-axis of a 96-well microtiter plate, leaving the last column as a blank (no added drug or bacteria). We evaluated all drug combinations of meropenem and the remaining four drugs (cefepime, gentamicin, ciprofloxacin and TMP/SMX). After 21 h of incubation at 36 °C in LB media (Bertani, 1951), we measured the OD_600_ in each well. Finally, we obtained isobolograms for the cefepime and meropenem combination and selected isolates using the BIGL library (Van der Borght et al., 2017) in the R platform (R Core Team, n.d.).

## Acknowledgements

This study was funded by the National Institutes of Health, (grant K08AI119182 from NIAID available to RJW) and the German Research Foundation, DFG (fellowship BA 6186/1-1 to CB).

## Supplementary figure legends

**Figure 4 – Supplementary Figure 1.**
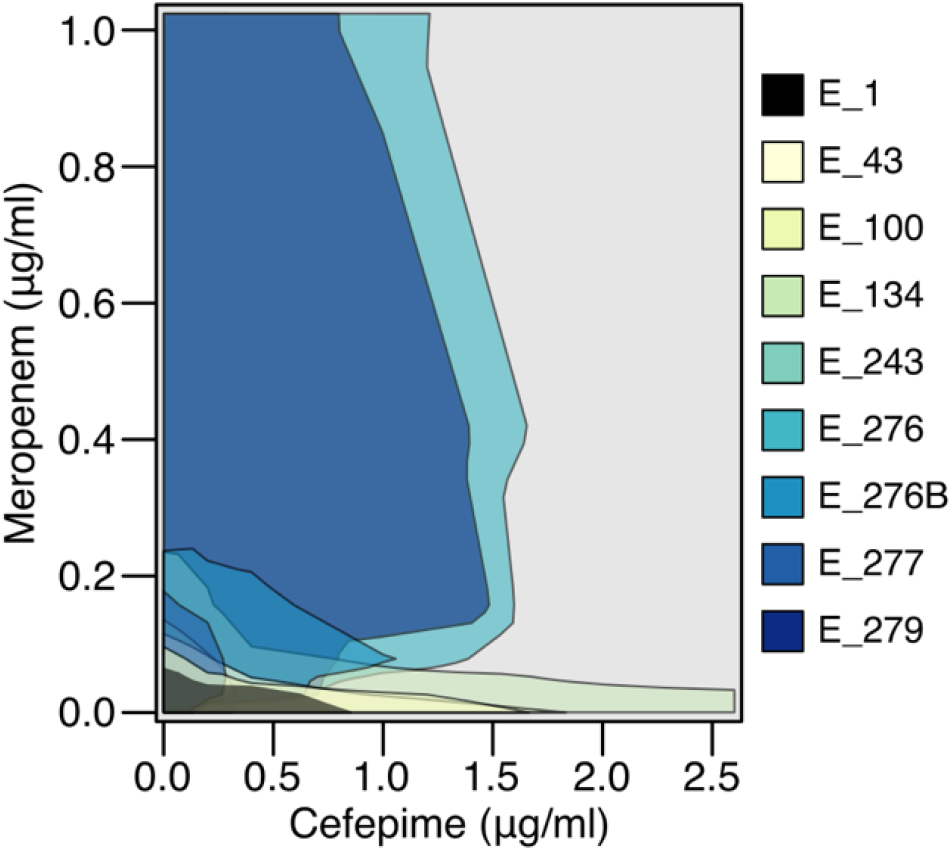
Isobologram curves for all isolates in a grid of concentrations of meropenem and cefepime inhibiting 50% of normal growth (IC50).

**Figure 4 – Supplementary Figure 2.**
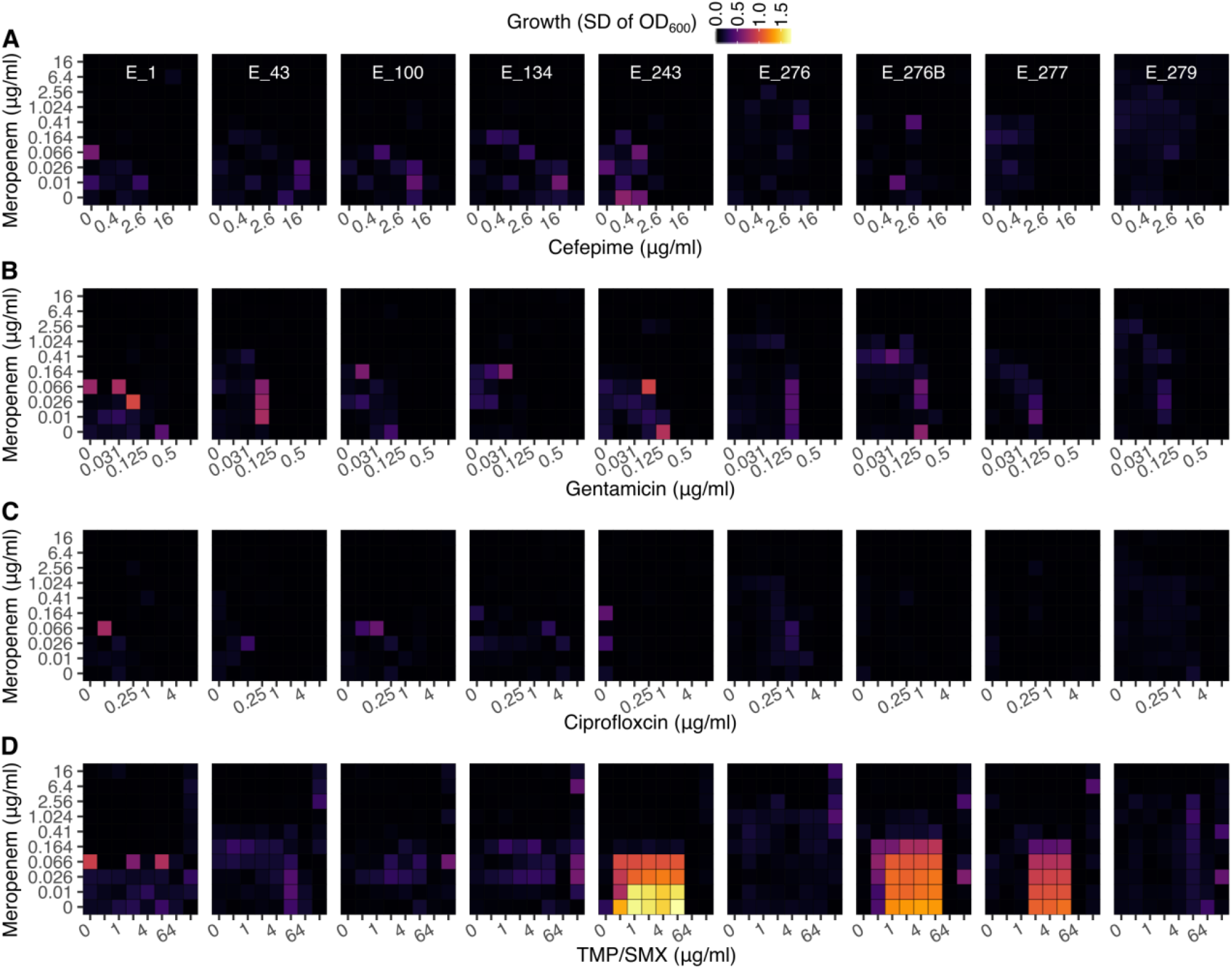
Standard deviation of two independent replicates across a grid of increasing concentrations of meropenem (Y-axis) and (A) cefepime, (B) gentamicin, (C) ciprofloxacin and (D) sulfamethoxazole/trimethoprim (TMP/SMX, or bactrim; X-axis) using the checkerboard assay.

## Source data legends

**Figure 2 – Source data 1 (separate file). Genomic variants identified within all isolates**.

**Figure 3 – Source data 1 (separate file). Growth measures**. Optical density (OD_600_) after 21h of growth from three independent replicates for each isolate and five different antibiotics.

**Figure 4 – Source data 2 (separate file). Checkerboards**. Optical density (OD_600_) after 21h of growth from two independent replicates in the presence of two drugs. We considered combinations of meropenem against four other antibiotics.

